# Assessing the limits of zero-shot foundation models in single-cell biology

**DOI:** 10.1101/2023.10.16.561085

**Authors:** Kasia Z. Kedzierska, Lorin Crawford, Ava P. Amini, Alex X. Lu

## Abstract

The advent and success of foundation models such as GPT has sparked growing interest in their application to single-cell biology. Models like Geneformer and scGPT have emerged with the promise of serving as versatile tools for this specialized field. However, the efficacy of these models, particularly in zero-shot settings where models are not fine-tuned but used without any further training, remains an open question, especially as practical constraints require useful models to function in settings that preclude fine-tuning (e.g., discovery settings where labels are not fully known). This paper presents a rigorous evaluation of the zero-shot performance of these proposed single-cell foundation models. We assess their utility in tasks such as cell type clustering and batch effect correction, and evaluate the generality of their pretraining objectives. Our results indicate that both Geneformer and scGPT exhibit limited reliability in zero-shot settings and often underperform compared to simpler methods. These findings serve as a cautionary note for the deployment of proposed single-cell foundation models and highlight the need for more focused research to realize their potential.^2^

## 1 Introduction

The emergence of foundation models in machine learning has been both transformative and rapid, as evidenced by the success of systems like ChatGPT [1] and DALL·E [2]. Foundation models are machine learning methods pretrained on huge amounts of data, where the aim of the pretraining is to enable models to capture universal patterns in data [3]. These models serve as adaptable starting points that can either be fine-tuned, which involves a small amount of additional training to prompt the model to produce specific predictive outputs, or used zero-shot, which involves extracting the model’s internal representation of input data (an “embedding”) for downstream analysis with no further task-specific training.

In single-cell biology, the foundation model framework offers an avenue for automating complex tasks, such as cell type identification and gene expression prediction. Emerging research has begun to explore the potential of foundation models in single-cell biology, particularly in single-cell transcriptomics, with several models now available. These include scBERT [4], Geneformer [5], scGPT [6], tGPT [7], scFoundation [8], SCimilarity [9], GET [10], GeneCompass [11], and GENEPT [12] which all present themselves as general models applicable to diverse analyses.

To evaluate their models, most previous works—including scGPT and Geneformer—rely on fine-tuning to specialize task-specific models. While this approach is a well-established practice in fields like natural language processing, its limitations become evident when applied to single-cell biology. Firstly, fine-tuning commonly requires a prediction problem with defined labels. However, much of the work in single-cell biology is inherently exploratory, where labels may not be available *a priori*. For instance, biologists often cluster latent representations of single-cell gene expressions to discover new cell types without pre-existing knowledge or imposed bias on the discovery process [13, 14, 15]. Secondly, the practicality of fine-tuning poses challenges for many labs. Even minimally fine-tuning foundation models can require extensive GPU resources, given that, for example, scGPT’s architecture relies on the use of FlashAttention [16] not available for older and smaller graphics cards^3^. Finally, zero-shot evaluation helps test the claim that pretraining promotes a foundational understanding of biology by exposing whether pre-training provides meaningful improvement over randomly initialized untrained models [17, 18, 19]. In alignment with these challenges, zero-shot capabilities have been rigorously evaluated in many other biological domains. In microscopy image analysis, for example, mainstream computer vision models have been shown to retrieve relevant image phenotypes without fine-tuning [20]. Similarly, language models tailored for protein sequences provide useful features for various protein engineering tasks even in zero-shot settings [21, 22, 23].

In this study, we assessed the zero-shot performance of two proposed foundation models in single-cell biology: Geneformer [5] and scGPT [6] - representative examples in a rapidly evolving field. Our assessment covers a range of tasks, including the utility of embeddings for cell type clustering, batch effect correction, and the effectiveness of the models’ input reconstruction based on the pretraining objectives (Fig. 1). Our findings indicate that Geneformer and scGPT are unreliable when applied in zero-shot scenarios. In tasks such as clustering and batch effect correction, they do not outperform simpler dimensionality reduction techniques. Further, our evaluation reveals that their pretraining objectives do not provide meaningful or useful information for biological applications. Together, our results caution against the use of proposed single-cell foundation models in zero-shot settings and suggest that current pretraining methods may not be initializing models with a general basis for transfer across biological settings.

**Figure 1:**
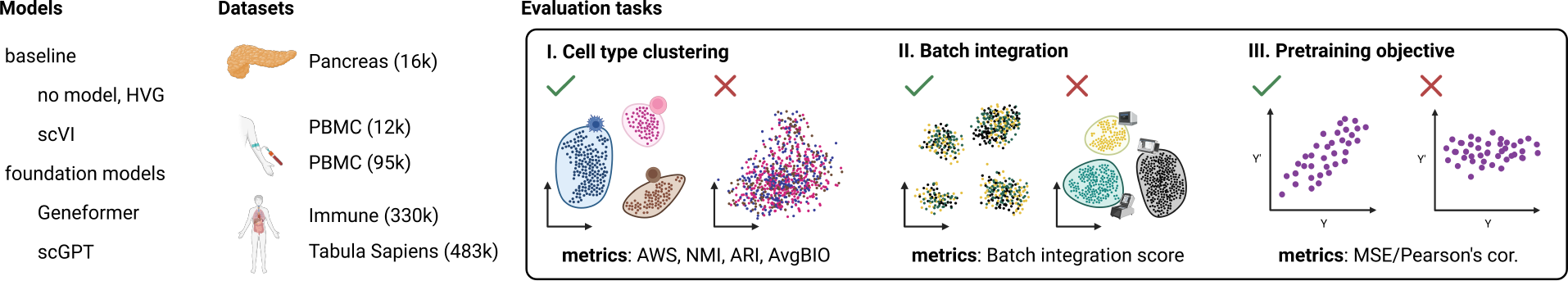
Overview of the evaluation setup. Our evaluation framework centers around two proposed foundation models, Geneformer and scGPT, and compares them to established methods like scVI and simpler strategies such as selecting highly variable genes (HVG) or predicting mean expression. To ensure comprehensive assessments, we curated a diverse set of five datasets. Our evaluation encompasses multiple facets, including the quality of cell embeddings for tasks like cell type clustering and batch integration. Additionally, we scrutinized the models’ performance with respect to their pretraining objectives.

## 2 Methodology

### 2.1 Models and baselines

We evaluated two proposed foundation models for single-cell transcriptomics: Geneformer [5] and scGPT [6]. We chose these models because they offer pretrained weights (whereas several other possible models did not have public weights at time of evaluation) and have been trained using unsupervised objectives on extensive datasets (ca. 30M single-cell transcriptomes). Here, we provide an overview of these models, including their practices for extracting cell embeddings, or latent representations of single-cells, which we follow for our analyses.

Both models accept single-cell gene expression vectors as input but represent input data differently. The input to the Geneformer model is a ranked list where the gene’s position represents the gene’s expression relative to the remaining genes in the cell. The model leverages a BERT-inspired architecture with 6 Transformer layers, each with 4 attention heads. Geneformer is trained using a modification of the masked language modeling (MLM) task, where the model is trained to recover randomly selected genes that are masked or corrupted. Since genes are ordered by their expression, this effectively predicts gene expression relative to other genes. The model outputs gene embeddings, which are subsequently decoded into gene predictions. A cell embedding is calculated by averaging over all gene embeddings extracted for that cell. Genefomer was pretrained on 27.4M human single-cell transcriptomes (excluding malignant and immortalized cells).

scGPT preprocesses each gene expression vector by independently binning values into 50 equidistant bins where the lowest bin is the lowest expression and the highest bin the highest expression. Next, the binned values and the gene token (i.e. a unique index for each gene) are separately embedded, and summed in the embedding space, jointly representing the gene and its binned expression. Like Geneformer, scGPT uses an MLM task. However, scGPT directly learns a cell embedding, which is integrated into its pretraining loss of predicting masked genes: scGPT first predicts a masked gene expression bin and a cell embedding from unmasked genes and then, in a second step, further iteratively refines masked gene expression using the cell embedding predicted in the first step. This means that scGPT outputs two sets of binned gene predictions in its pretraining task, first from unmasked genes alone and second from conditioning on the cell embedding. In our effort to understand the generalization of the pretraining objectives, we analyzed both. Finally, compared to Geneformer, scGPT has 3× the parameters, using 12 Transformer layers with 8 attention heads. scGPT is available in several variants, pretrained on multiple different datasets. In our analyses, we focused on three variants of scGPT: pretrained on 814,000 kidney cells (scGPT kidney), on 10.3 million blood and bone marrow cells (scGPT blood), and on 33 million non-cancerous human cells (scGPT human).

For baselines in evaluating cell embeddings, we compared Geneformer and scGPT against selecting highly variable genes (HVGs). We standardize to 2,000 HVGs across all experiments. In addition, we compared all methods to scVI, a scalable generative model [24] which we trained on each individual dataset. While this means that we deploy scGPT and Geneformer zero-shot while training scVI on target data unsupervised, we reasoned this set-up reflects practical settings where resources are available to train lightweight models, but not to fine-tune large models. For the evaluation of the pretraining objective, we used the mean estimates or average ranking as a reference.

### 2.2 Datasets

To assess the quality of cell embeddings and performance on batch integration tasks, we used five distinct human tissue datasets (Table 1). These datasets include samples from the pancreas [25], two sets of peripheral blood mononuclear cells (PBMCs) [26, 27], a cross-tissue immune cell atlas [28], and a multi-organ human cell atlas [29]. Each dataset poses unique challenges relevant to single-cell analysis, such as the distinction between well-defined and less well-defined cell type clusters, the integration of different technical batches within the same tissue, and the unification of data across multiple tissues.

**Table 1:** Overview of the used datasets.

| Dataset name | Description | No. of cells | No. of labels | No. of batches | Ref. |
| --- | --- | --- | --- | --- | --- |
| Pancreas | Cells from human pancreas created by combining data spanning 5 studies. | 16k | 14 | 6 | [25] |
| PBMC | PBMCs from a healthy donor. | 12k | 9 | 2 | [26] <sup>4</sup> |
| PBMC | PBMCs from a healthy donor. | 95k | 10 | 1 | [27] |
| Immune | Immune cells extracted from 16 different tissues across 12 adult organ donors. | 330k | 45 | 31 | [28] |
| Tabula Sapiens | Cells from 24 different tissues across 15 human donors. | 483k | 177 | 27 | [29] |

Among the selected datasets, the Pancreas dataset partially overlapped with the data used to pretrain Gene-former. We conducted evaluations using both the complete Pancreas dataset and its non-overlapping subset. The results were highly consistent between the two, leading us to include the entire Pancreas dataset for simplicity in this evaluation. At the time of dataset selection, information on the data used for scGPT’s pretraining was unavailable, preventing us from determining any potential overlaps at the time of our evaluations.

### 2.3 Evaluation metrics

In this work, we evaluated the cell embedding space for its ability to separate known cell types correctly and to integrate different batches. We also evaluated the performance of the models at the pretraining task by evaluating their reconstruction accuracy.

#### 2.3.1 Average silhouette width (ASW) and average BIO (AvgBIO) scores

One key aspect of evaluating cell embeddings is the degree to which cell types are distinct within the embedding space. To assess this, we employ metrics based on the Average Silhouette Width (ASW) [25] and the Average Bio (AvgBIO) scores [6]. Briefly, ASW is computed by taking the difference of the between-cluster and within-cluster distances and dividing this by the larger of the two values. ASW is normalized to a range between 0 and 1, where 0 signifies strong within-cluster cohesion, 0.5 indicates overlapping clusters, and 1 denotes well-separated clusters. Higher ASW indicates better performance in separating clusters. AvgBIO is the arithmetic mean of three individual metrics: ASW, Normalized Mutual Information (NMI), and Adjusted Rand Index (ARI), as defined in [6]. NMI and ARI are calculated based on Louvain clusters generated directly from the embedding space [25, 6]. AvgBIO is normalized to a 0-1 scale, with higher values indicating better alignment between clusters and ground truth labels.

#### 2.3.2 Batch integration score

To evaluate batch integration, we used a variation of the AWS score (as described in [25]). Briefly, the silhouette scores are calculated with respect to the batch label by taking only its absolute value, where a score of 0 is equivalent to absolute mixing and any deviation from 0 indicates the presence of a batch effect. To keep with the used convention, the score is then subtracted from 1, resulting in final scores on a scale between 0 and 1, where a final score of 0 suggests complete separation of the batches and strong batch effect and 1 signifies a perfect batch mixing and integration.

#### 2.3.3 Reconstructing gene expression

To evaluate the performance of scGPT in its pretraining objective, we used the mean squared error (MSE), as used by the original authors for the model’s loss [6]. For reference, we calculated the mean bin of a gene from the dataset and calculated the MSE for the input bins and the mean bins.

To evaluate Geneformer’s performance in its pretraining objective, we measured Pearson’s correlation between the true and predicted rankings. For that, we transformed ordered outputs into scaled (from 0 to 1) rankings, where the highest expressed genes were assigned a rank of 1. Geneformer can output a sequence of up to 2,048 genes. To handle variable-length input sequences during batch processing, padding is applied to match the length of the longest sequence in the batch. In our evaluations, we truncated the model’s output to match the input sequence length. As a baseline for comparison, we determined the mean ranking position of each gene based on the rankings with that gene present. We then computed Pearson’s correlation coefficient between the input ranking and the ranking composed of mean positions for those genes.

## 3 Results

### 3.1 Cell type clustering

Current proposed single-cell foundation models produce cell embeddings. These embeddings are intended to project potentially noisy gene expression measurements to a more biologically relevant latent space and to thus improve our ability to resolve cell types, consistent with previous machine learning methods in this field (including scVI) [30, 31, 24, 32]. Both scGPT and Geneformer fine-tune their cell embeddings for cell type classification. However, this strategy fails in more exploratory contexts where cell composition in the dataset may not be known. For these applications, foundation models must produce robust cell embeddings zero-shot. Therefore, we evaluated the zero-shot performance of scGPT and Geneformer in separating known cell types across multiple datasets. We also compared these approaches to a baseline strategy of selecting highly variable genes (HVGs) and to an unsupervised learning model, scVI.

We evaluated cell type clustering using two metrics, ASW and AvgBIO. For both metrics, Geneformer and scGPT performed worse than our baseline strategies. For ASW, scVI consistently performed well, achieving a median ASW of 0.54 and hitting a low of 0.49 in the Tabula Sapiens dataset (Fig. 2A). Geneformer’s performance was more variable, with scores ranging from a high of 0.52 in the PBMC (95k) dataset to a low of 0.38 in the Pancreas (16k) dataset. scGPT’s performance was comparable with scVI, with the median ASW of the former equal to 0.53. scGPT outperformed scVI for blood datasets but fell behind for Pancreas and Tabula Sapiens datasets. Notably, HVG outperforms Geneformer in all datasets except PBMC.

**Figure 2:**
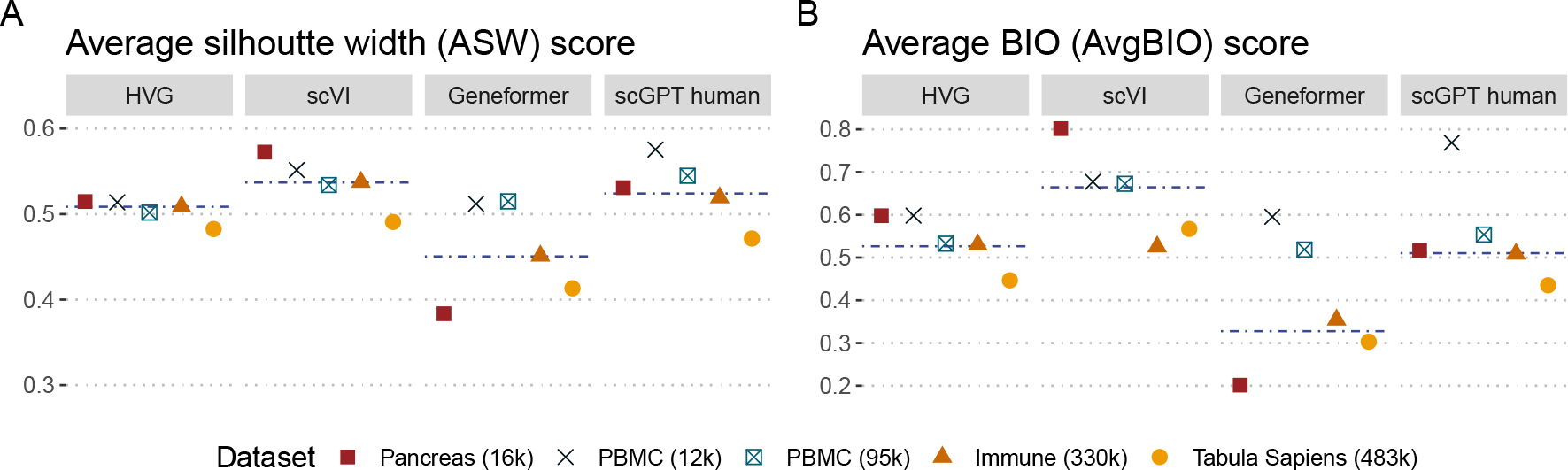
Proposed single-cell foundation models fail to outperform cell embeddings derived from HVG or generated using the scVI model. **A** Average silhouette width score and **B** Average BIO score (described in Section 2.3.1) calculated on the highly variable genes (HVG) of the log normalized input data and on the embeddings extracted from scVI, scGPT, and Geneformer models. Median value annotated with a dashed line. A higher score indicates better performance in separating clusters.

For AvgBIO, scVI similarly surpassed all other models in AvgBIO score in three out of five datasets (Fig. 2B). Notably, while scGPT performs exceptionally well on the PBMC (12k) dataset, registering an AvgBIO score of 0.77, it shows inconsistent performance across datasets, with other datasets decreasing in performance relative to HVG (dropping from 0.60 to 0.52 on the Pancreas dataset). In contrast, scVI consistently had equivalent or better performance than HVG.

Foundation models usually employ self-supervised tasks to enable scalability since they can train on any dataset, not just ones with labels [3]. However, it is unclear if pretraining on larger datasets improves the cell embeddings learned by proposed single-cell foundation models. Therefore, we next assessed the impact of the pretraining dataset on model performance. We focused on scGPT due to its release of weights pretrained on various datasets. We assessed four different models: randomly initialized scGPT as a baseline with no pretraining, scGPT pretrained on 814,000 kidney cells (scGPT kidney), on 10.3 million blood and bone marrow cells (scGPT blood), and on 33 million non-cancerous human cells (scGPT human). One limitation of our analysis is the smaller models are trained on tissue-specific data, confounding if differences in performance are due to size or the composition of dataset. However, at minimum, scGPT human includes all data used to train scGPT blood and scGPT kidney.

Our analysis indicates that pretraining does improve cell-type clustering, with the median score for all models improving over random models (Figure 3, Table S1).

**Figure 3:**
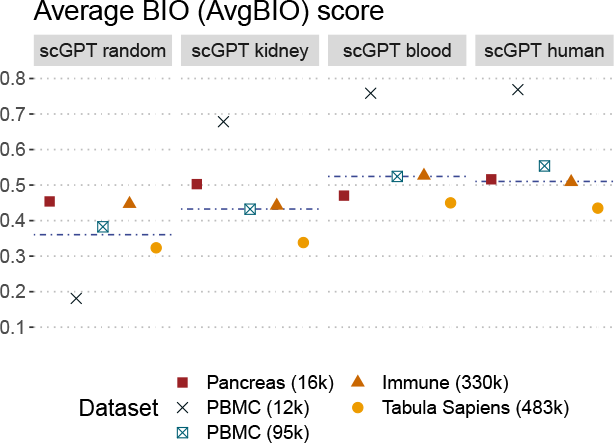
Size of the pretraining dataset correlates with the performance at separating the cell types in cell embedding space. Average BIO score (described in 2.3.1) calculated on the embeddings extracted from selected variations of the scGPT models. The dashed line marks the median score across datasets.

Overall, our findings demonstrate that foundation models in zero-shot configurations perform inconsistently compared to cell embeddings derived from HVG or generated using the scVI model. Evaluating variants of the scGPT model highlights that while pretraining confers some benefit, past a certain limit, larger and more diverse datasets fail to confer further benefits.

### 3.2 Batch integration

Next, we sought to assess the zero-shot capabilities of proposed single-cell foundation models in batch integration. Single-cell transcriptomics experiments, like all biological experiments, are impacted by batch effects - systematic technical differences present when integrating data over different experiments, sequencing technologies, or even when the experiment is reproduced for the same biological replicates. Due to batch effects, tasks like mapping a new experiment to a reference atlas to identify the cell types present in the data can fail. Hence, a common task in single-cell analysis is to eliminate batch effects without removing meaningful biological differences, allowing for data integration [13, 14, 15].

We began with a qualitative evaluation of the Pancreas dataset, a common batch integration benchmark that includes data from five different sources [25]. As commonly done in single-cell transcriptomics, we used UMAP projections to visually inspect embeddings (Fig. 4). By annotating the UMAP by cell type (Fig. 4A) versus experimental technique (Fig. 4B), we jointly assess if cell embeddings correct for batch effects stemming from techniques while still retaining cell type identity. As demonstrated by the UMAP of all genes, batch effects impact this data, with experimental techniques separated (and also forming sub-clusters, some of which are a result of different batches taken with the same technique) (Fig. 4B).

**Figure 4:**
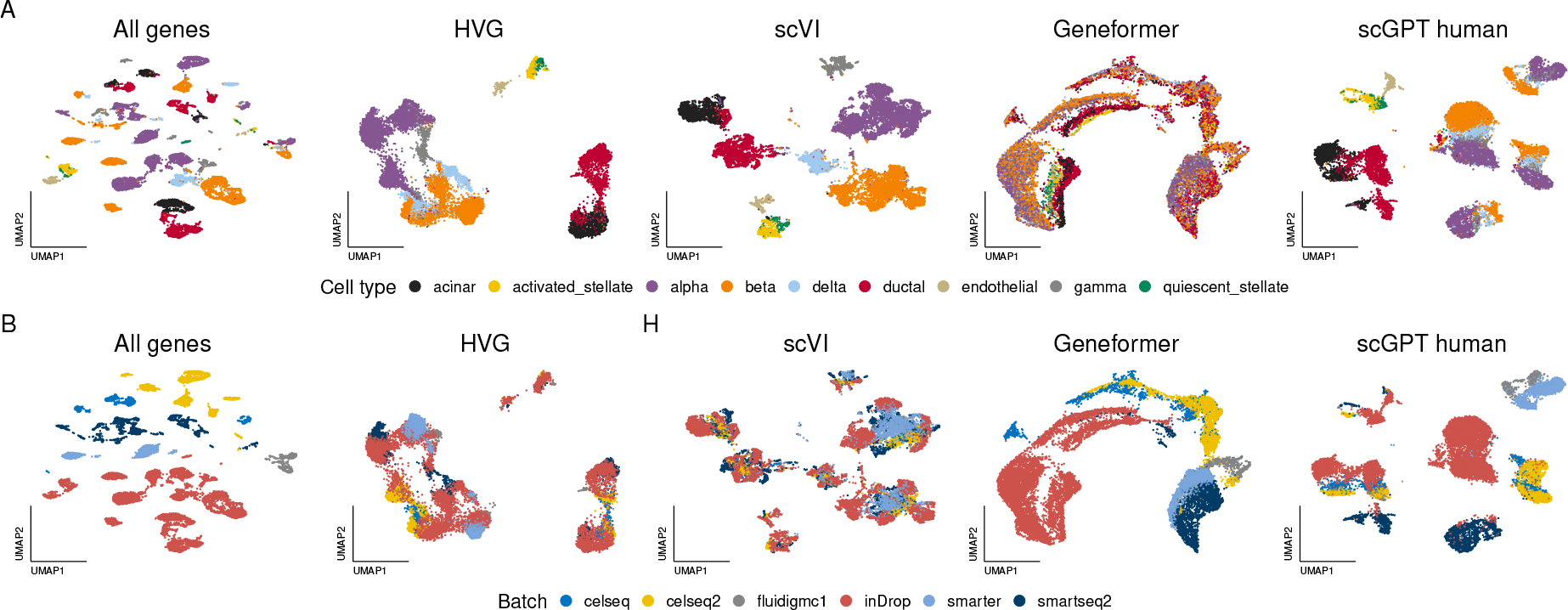
Zero-shot foundation models perform poorly at integrating batches of Pancreas dataset. **A, B** Visualization of the UMAP projections of the pancreas dataset using normalized input data, normalized input data preselected for highly variable genes (HVG), and cell embeddings generated by scVI, Geneformer and scGPT human. Cells are color-coded by cell type (A) and batch (B).

**Figure 5:**
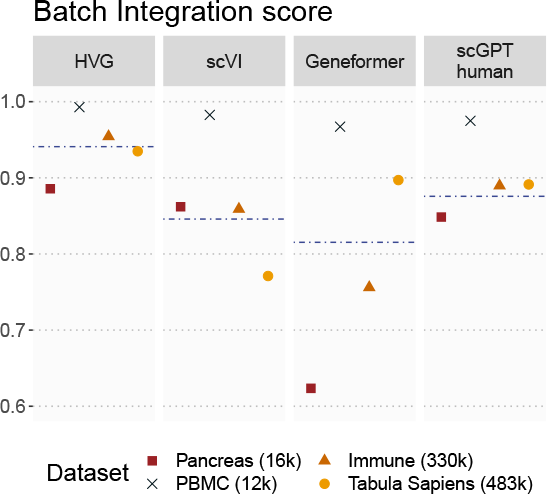
HVG selection outperforms proposed foundation models. Batch integration score (described in Section 2.3.2) calculated for all four datasets with at least two batches.

Overall, we observed that while Geneformer and scGPT-human can integrate different experiments conducted with the same experimental technique, they generally fail to correct for batch effects between techniques. As depicted in Fig. 4A, the cell embedding space generated by Geneformer fails to retain information about cell type, and any clustering is primarily driven by batch effects (Fig. 4B). On the other hand, the space created by scGPT offers some separation of cell types (Fig. 4A), but the primary structure in the dimensionality reduction is driven by batch effects (Fig. 4B). In contrast, even the simple baseline of selecting highly variable genes (HVG) qualitatively produces a similar or better result to scGPT, with the Smarter technique now being integrated with InDrop. Finally, we observed that scVI mostly integrates this dataset, forming clusters primarily due to cell type, with most techniques in the same cluster.

To support these qualitative results, we produced batch integration metrics for each of our four datasets with more than one batch (Fig. 4C). Geneformer underperforms compared to both scGPT and scVI across most datasets, achieving a median batch integration score of only 0.8. scVI outperforms scGPT in datasets where the batch is restricted to the technical variation (Pancreas and PBMC datasets), and scGPT performs better in more complex datasets where both technical and biological batch effects are present (Immune and Tabula Sapiens datasets). Surprisingly, the best batch integration scores for all datasets were achieved by selecting HVG. This observation is slightly different from our qualitative evaluations of the UMAPs where scVI performs better, and can be explained by shifts in our rankings calculating metrics in full rather reduced dimensions as seen in Fig. S2 (we note that trained proposed foundation models underperform baselines in both settings).

In summary, our evaluation suggests that Geneformer and scGPT are not fully robust to batch effects in zero-shot settings, often lagging behind existing methods like scVI, or simple data curation strategies like selecting for HVG, particularly when batch effects are more severe.

### 3.3 Pretraining objective

Next, to understand why Geneformer and scGPT underperform compared to baselines zero-shot, we posited two hypotheses. First, it could be that the masked language modeling pretraining framework used by both scGPT and Geneformer does not produce useful cell embeddings. The second could be that scGPT and Geneformer have failed to generalize the pretraining task. Understanding this distinction could produce insights for future directions. For example, if the models are reconstructing masked gene expression well for our evaluation datasets but still failing to produce informative cell embeddings, this implies that a different task may need to be designed; while if the models fail to predict gene expression accurately, improvements to learning the pretraining task could still potentially improve the cell embeddings of these models.

Evaluating whether models reconstruct masked gene expression accurately requires us to select how many genes are masked in input. In training, both models select a percentage of genes to mask. However, following a similar procedure for evaluation introduces stochasticity, and re-running random samples and/or iterating over genes to account for this is computationally expensive. We, therefore, use *all* genes unmasked as input. Not only does this eliminate stochasticity from sampling masked genes, but it also reflects the maximally informative setting where models are asked to reconstruct genes given complete, not partial, input.

To gauge the quality of these reconstructions, we compared them to their true values. For scGPT, we compared the bin value for each gene. Since scGPT produces gene predictions at two stages (with and without conditioning from its cell embedding), we report both. For Geneformer, we compared the gene rankings. scGPT largely fails to reconstruct gene expression, especially in its gene expression prediction (GEP), i.e. under the MLM objective, while Geneformer shows better agreement between the prediction and the true value (Fig. 6). Without conditioning on cell embedding, scGPT predicts the mean value of the input bin across all bin values (Fig. 6A). When conditioned on cell embeddings (GEPC), performance improves for the Immune (330k) dataset (Fig. 6B) but remains inconsistent across various datasets (Fig. S3, Table S4).

**Figure 6:**
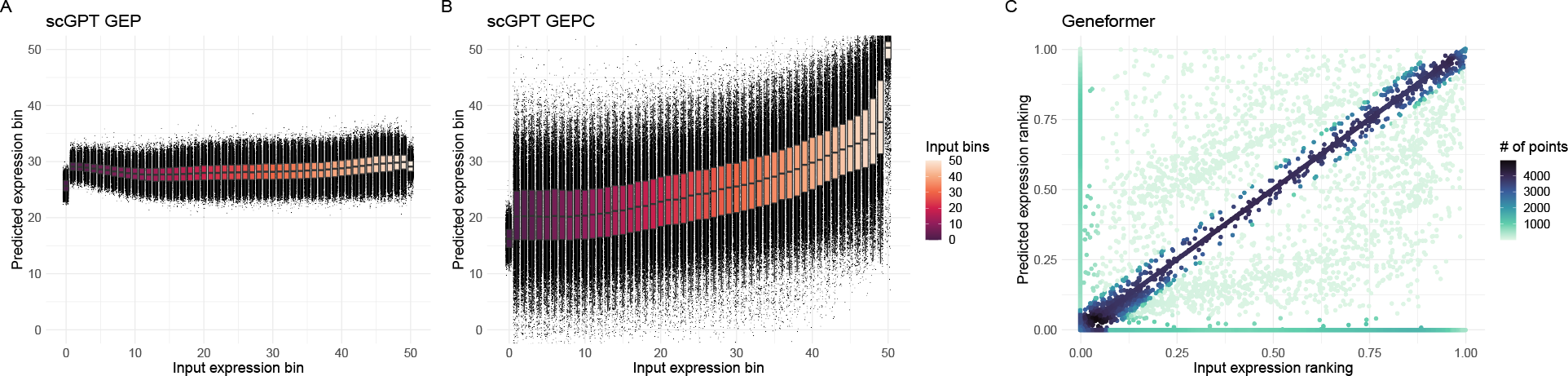
scGPT struggles with reconstructing gene expression. Reconstruction of the Immune (330k) dataset with **A** the output of the MLM task in scGPT model, **B** expression prediction from cell embedding in scGPT, and **C** output of masked prediction in Geneformer model (color signifies the number of points represented by one dot).

As a baseline to confirm this behavior, we compared the performance of scGPT against a naive baseline of just predicting the mean expression value of a gene. Surprisingly, this baseline prediction outperformed all scGPT variants when not using cell embeddings (Fig. 7A, Table S3), with only marginal improvements observed when conditioning on cell embedding.

**Figure 7:**
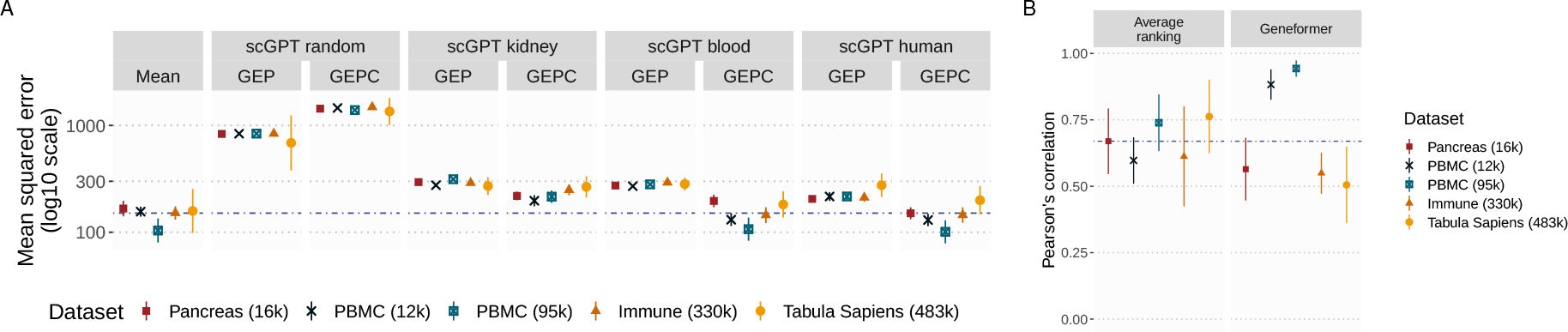
scGPT and Geneformer perform similarly to when mean values or average ranking are used. **A** MSE for the reconstructed input compared to the ground truth for the two objectives of scGPT: masked expression prediction (gene expression prediction, GEP) and gene expression prediction from cell embedding (GEPC). Mean and standard deviation ranges shown as point and full lines, respectively. **B** Pearson’s correlation between the input ranking and the average ranking and the output of Geneformer. Data shown as a range with a point indicating mean value and lines annotating the standard deviation range for all datasets. The median value of the MSE calculated for the average bin value or for the correlation of the average ranking across all datasets indicated by a dashed line. Mean and standard deviation ranges shown as point and full lines, respectively.

Geneformer does not directly predict expression but generates a ranked list of genes. To evaluate Geneformer, we therefore measure Pearson’s correlation between the predicted ranking of genes and the actual gene ranking. Under its MLM objective, Geneformer predicts the most likely gene at a given position. There is a moderate correlation between the output and input rankings for the Immune dataset (mean Pearson’s correlation of 0.55, Fig. 6C). While there is a good agreement among a large number of genes, the model fails to predict some genes while also predicting genes not present in the input ranking. Geneformer outperforms the average ranking in the blood datasets, while achieving lower scores for the remaining datasets (Fig. 7B, Fig. S3, Table S4)

## 4 Discussion

In this work, we evaluated two proposed foundation models for single-cell biology – Geneformer and scGPT – and demonstrated their unreliability in zero-shot settings. In cell type clustering analyses, both models fail to improve reliably over scVI. Critically, for some datasets, the proposed foundation models perform worse at clustering cell types than just selecting highly variable genes. At least for scGPT, while we demonstrated that pretraining at least confers some benefit, models trained on larger and more diverse datasets may not do better than models trained on data from a single tissue of a reasonable scale.

We also demonstrate that these models are not fully robust to batch effects in zero-shot settings, often lagging behind methods like scVI or simple data curation strategies like selecting for HVG.

Together, our results caution against using current single-cell transcriptomic foundation models in zero-shot settings. Our analyses provide some insight on where future work needs to be concentrated to build bonafide foundation models that are truly useful in these settings. Intriguingly, we found no relationship between a model’s ability to solve their masked language modeling (MLM) task and zero-shot performance: while Geneformer is capable of reconstructing input gene rankings closely, and while scGPT primarily defaults to predicting the average gene expression bin, scGPT outperformed Geneformer in all evaluations. Our work suggests that greater research must be dedicated to articulating the relationship between pre-training task, pre-training dataset, and performance on downstream analysis tasks.

## Footnotes

2 The code used for our analyses can be accessed at https://github.com/microsoft/zero-shot-scfoundation.

3 FlashAttention currently supports Ampere, Ada, or Hopper GPUs (e.g., A100, RTX 3090, RTX 4090, H100) and Turing GPUs (T4, RTX 2080). Currently, no plans exist to support other GPUs, such as the popular V100.

4 Data available via data.pbmc_dataset function from scvi-tools [26] Python package.

