## Supplemental Figures for "Assessing the limits of zero-shot foundation models in single-cell biology"

### 5 Supplementary Material

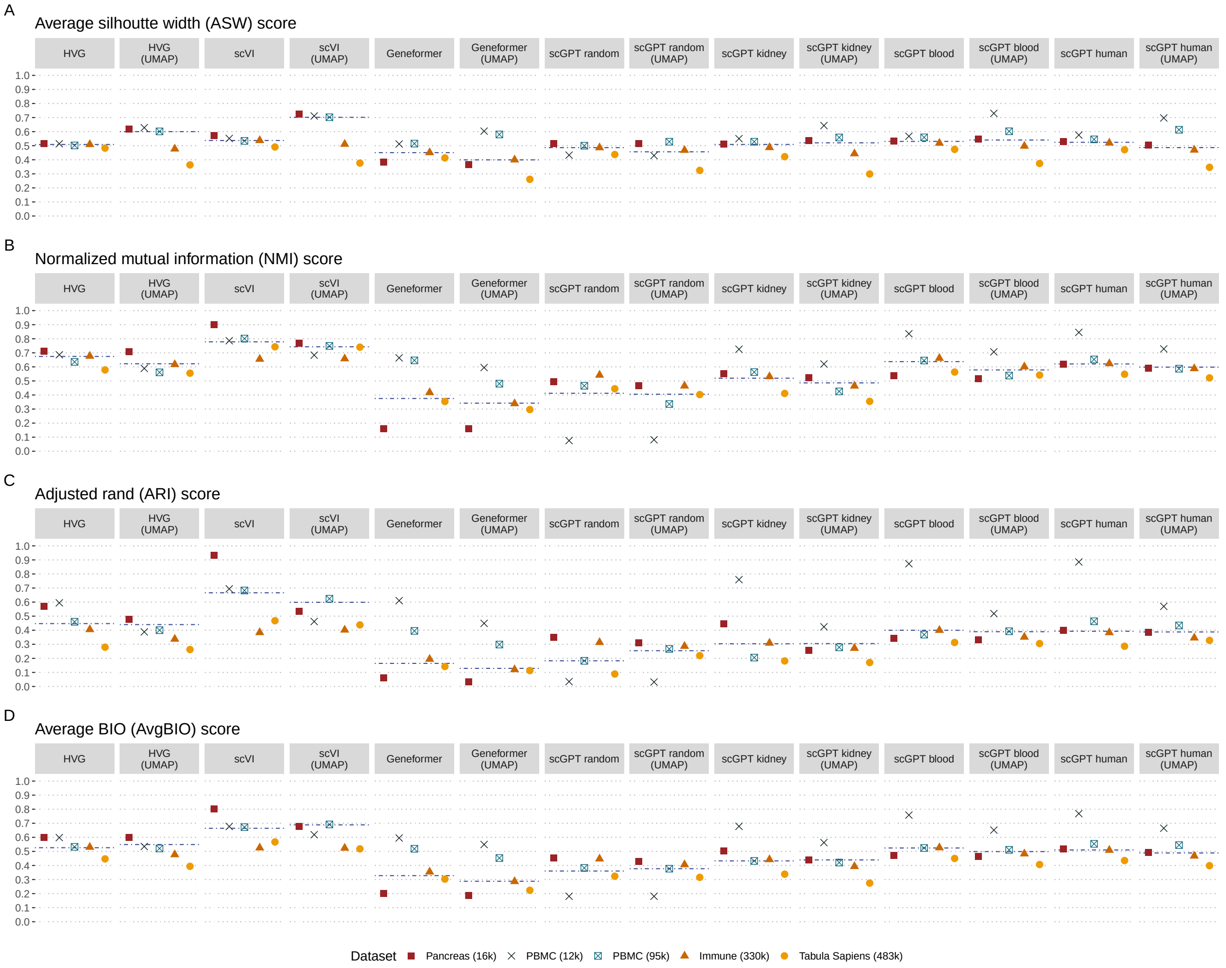


Figure S1: **UMAP projection of the embeddings improves cell type separation for scVI.** Scores used for assessing cell types separation in cell embedding space across raw and UMAP projected embeddings. **A** Average silhouette width (ASW) score, **B** Normalized mutual information (NMI) score, **C** Adjusted rand (ARI) score and **D** Average BIO (AvgBIO) score - an average of the other scores.

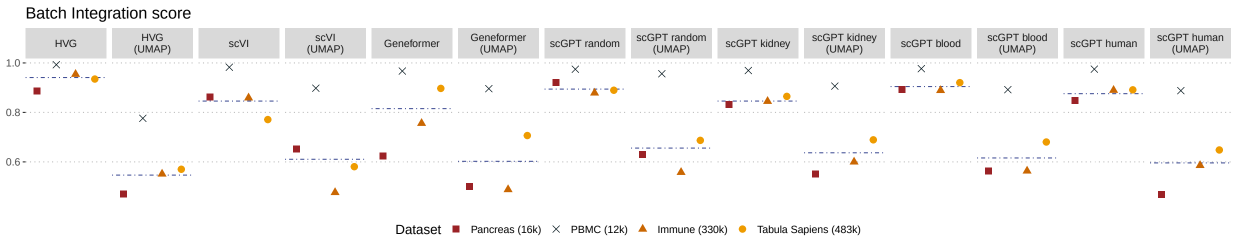


Figure S2: **UMAP projection of the embeddings results in lower batch integration scores.**

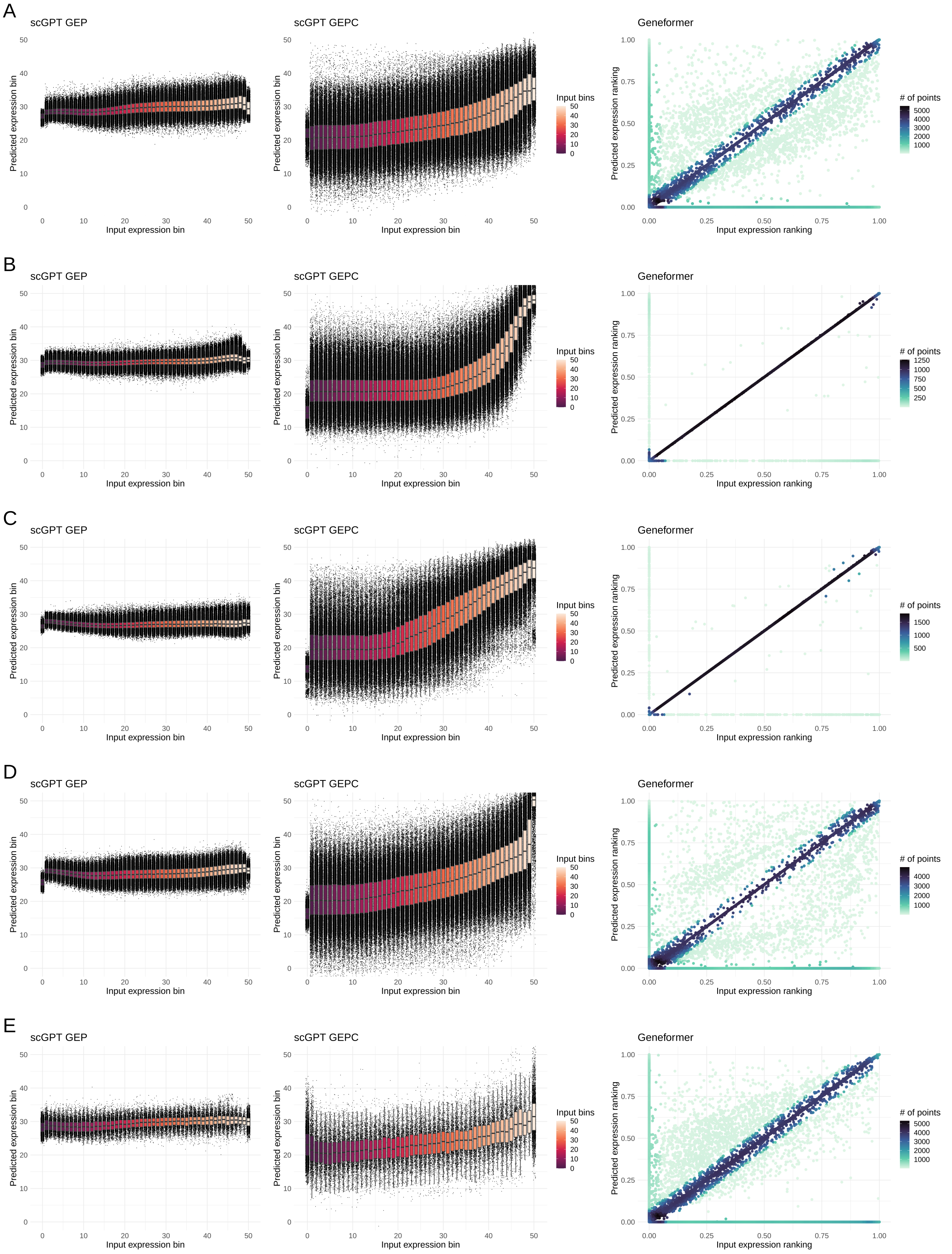


Figure S3: **Performance of the proposed foundation models with respect to reconstructed expression bin (scGPT human) or ranking (Geneformer).** The predicted bins as a function of input bins for scGPT GEP (masked language modeling objective, left panel), scGPT GEPC (conditional on cell embedding, middle panel) and the agreement between input and output rankings for Geneformer (right panel) for **A** Pancreas, **B** PBMC (12k), **C** PBMC (95k) **D** Immune (330k) **E** Tabula Sapiens (483k) datasets shown.
